# The APOBEC3A deaminase drives episodic mutagenesis in cancer cells

**DOI:** 10.1101/2021.02.14.431145

**Authors:** Mia Petljak, Kevan Chu, Alexandra Dananberg, Erik N. Bergstrom, Patrick von Morgen, Ludmil B. Alexandrov, Michael R. Stratton, John Maciejowski

## Abstract

The APOBEC3 family of cytidine deaminases is widely speculated to be a major source of somatic mutations in cancer^1–3^. However, causal links between APOBEC3 enzymes and mutations in human cancer cells have not been established. The identity of the APOBEC3 paralog(s) that may act as prime drivers of mutagenesis and the mechanisms underlying different APOBEC3-associated mutational signatures are unknown. To directly investigate the roles of APOBEC3 enzymes in cancer mutagenesis, candidate *APOBEC3* genes were deleted from cancer cell lines recently found to naturally generate APOBEC3-associated mutations in episodic bursts^4^. Deletion of the *APOBEC3A* paralog severely diminished the acquisition of mutations of speculative APOBEC3 origins in breast cancer and lymphoma cell lines. APOBEC3 mutational burdens were undiminished in *APOBEC3B* knockout cell lines. *APOBEC3A* deletion reduced the appearance of the clustered mutation types *kataegis* and *omikli*, which are frequently found in cancer genomes. The uracil glycosylase UNG and the translesion polymerase REV1 were found to play critical roles in the generation of mutations induced by APOBEC3A. These data represent the first evidence for a long-postulated hypothesis that APOBEC3 deaminases generate prevalent clustered and non-clustered mutational signatures in human cancer cells, identify APOBEC3A as a driver of episodic mutational bursts, and dissect the roles of the relevant enzymes in generating the associated mutations in breast cancer and B cell lymphoma cell lines.

## MAIN

Early investigations into the patterns of somatic mutations in cancer genomes have revealed that both non-clustered and clustered mutations at cytosine bases commonly present at TCN (where N is any base) trinucleotide sequence contexts^1,2^. Previously recognized sequence preferences of the APOBEC3 family of cytidine deaminases, which target DNA and RNA of viruses and retroelements as part of the innate immune defense, led to the proposal that such mutations may represent APOBEC3 off-target activity^1,2^. Subsequent mathematical deconvolution of somatic mutational patterns across thousands of human cancer genomes led to the identification of APOBEC3-associated mutational signatures in more than 78% of cancer types and 56% of all cancer genomes analyzed to date, with a particular prominence in breast, bladder, and other cancer types^5,6^. Two mutational signatures of single base substitutions (SBS), termed ‘SBS2’ and ‘SBS13’, have been proposed to be caused by off-target APOBEC3 activities^5^.

The APOBEC3 hypothesis (Fig. 1a) proposes that one of the five APOBEC3 enzymes with a preference for TCN motifs deaminates cytosine bases in TCN motifs in single-stranded DNA (ssDNA)^3,7^. Subsequent processing of the resulting uracil base likely determines the type of mutation. Replication across the uracil bases is assumed to give rise to C>T mutations and thus possibly SBS2. Uracil excision by a glycosylase, such as UNG or SMUG1, and downstream processing by base-excision repair (BER) and translesion polymerases may give rise to C>T, C>G and C>A mutations and thus a combination of SBS2 and SBS13^3,7^. Consistent with this proposal, overexpression of individual human APOBEC3 enzymes in yeast and other models can result in SBS2 and SBS13-like mutations^8,9^.

**Figure 1.**
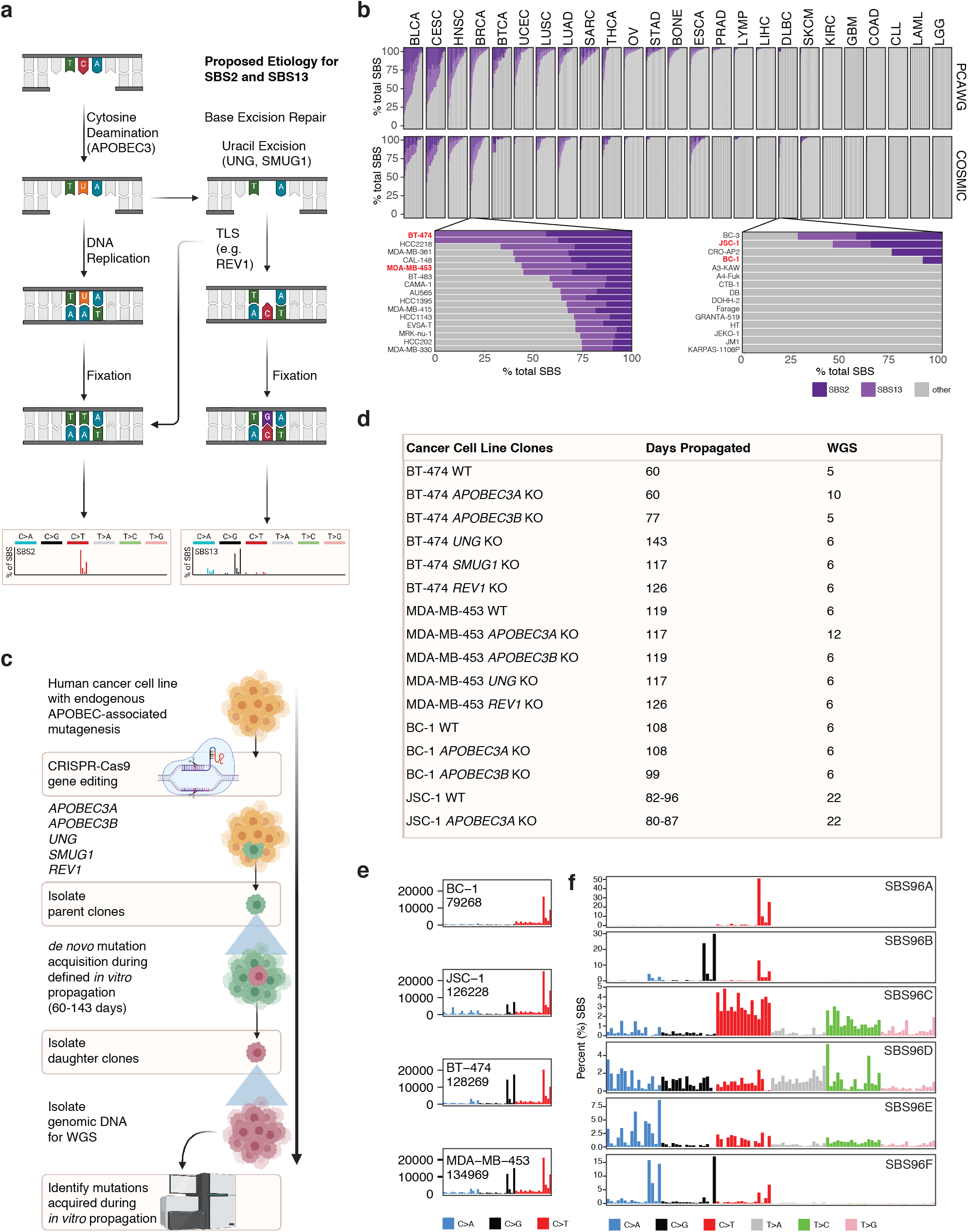
Using human cancer cell lines to investigate origins of APOBEC3-associated mutagenesis. **a)** Speculative mechanisms of APOBEC3-associated SBS2 and SBS13 mutational signatures in cancer. **b)** Prevalence of SBS2 and SBS13 in sequences from 780 COSMIC cancer cell lines (top panel) and 1,843 sequences from human cancers (bottom panel). Each bar represents a percentage of mutations attributed to the indicated mutational signatures in an individual cell line or a cancer sample from cancer types indicated on top (abbreviations in Table S1). BRCA and DLBC datasets are magnified to show individual cell lines including those chosen for further study highlighted in red. **c)** Experimental design used to track mutation acquisition over controlled *in vitro* timeframes. Following CRISPR-Cas9 targeting of candidate genes, single cells were isolated, grown into ‘parent clones’ and propagated in culture for 60-143 days. Following this period, individual cells were isolated from each parent population and grown into ‘daughter’ clones that were expanded for DNA isolation. DNAs from parent and daughter clones were subjected to WGS and mutations were identified in each clone. Subtraction of mutations identified in parent clones, from mutations present in their relevant daughters, reveals mutations acquired during the *in vitro* timeframes spanning the two cloning events. **d)** Sample overview. Numbers of days spanning the two subcloning events during which mutational acquisition was tracked were denoted under ‘Days Propagated’ and the total number of wild-type and knockout parent and daughter clones subject to sequencing is under ‘WGS.’ **e)** Cancer cell lines carry signatures of historic APOBEC3-associated exposures. Mutational profiles from individual cell lines are displayed according to the number (y-axis) of genome-wide 96-substitution classes denoted on horizontal axis, which are defined by the six color-coded SBS types and 16 possible alphabetically ordered trinucleotide sequence contexts at which each mutation type presents (order of individual substitutions follows standard format, detailed in Extended Fig. 4). **f)** Profiles of mutational signatures extracted *de novo* from 815,923 SBS identified across mutational catalogues of 4 stock cell lines and 136 parent and daughter clones. SBS (single base substitution), TLS (translesion synthesis), PCAWG (Pan-Cancer Analysis of Whole Genomes), WGS (whole-genome sequencing). Each signature is displayed according to the percentage (y-axis) of genome-wide 96-substitution classes denoted on horizontal axis, which follow standard representation (details in Extended Fig. 4).

Speculations regarding the contributions of endogenous APOBEC3 enzymes to mutations in human cancer cells and involvement of the subsequent DNA repair and replication mechanisms are supported by association-based studies, but not causal links^3,10,11^. Expression of both APOBEC3A and APOBEC3B correlates with the APOBEC3-associated mutational burdens in many cancers, albeit weakly^12–16^. Progress in testing the APOBEC3 hypothesis in a more natural setting has been hindered by differences between the human and murine APOBEC3 loci and the lack of human cancer cell models. As a result, there has been substantial debate regarding whether APOBEC3A, APOBEC3B, or other APOBEC3 enzyme(s) generate the majority of mutations seen in cancer^8,12,13,17–19^. High expression levels of APOBEC3B support a model in which APOBEC3B generates most APOBEC-associated mutations seen in cancer^12,13^. Further suggesting a potential mutator role, APOBEC3B is the major source of cytidine deaminase activity in breast cancer cell lines^12,13^. However, cancers that develop in carriers of a germline deletion of *APOBEC3B* often exhibit higher burdens of the relevant mutations suggesting a potential mutator role for additional APOBEC3 enzymes, at least in certain contexts^17,20^. Indeed, other correlative studies nominate APOBEC3A. APOBEC-associated mutations in cancer mostly present in a sequence context preferred by APOBEC3A^8^ and APOBEC3A was recently reported to have a stronger deamination activity compared to APOBEC3B in breast cancer cell lines^15^.

It is critical to establish whether APOBEC3 activity causes mutations in human cancer and to identify the relevant mutator paralog(s) in order to pursue proposed therapeutic strategies based on modulating APOBEC3 activities in cancer^21–28^ and to conduct future research into the unknown instigators of the speculative, mutagenic APOBEC3 behavior. Here, by CRISPR-Cas9 deleting the candidate APOBEC3 mutators from cancer cell lines that generate the relevant mutations naturally over time^4^, we provide the first experimental evidence in human cancer cells for a hypothesis put forward almost two decades ago^29^. Despite its minimal expression relative to APOBEC3B, we identify APOBEC3A as the major driver of episodic mutational bursts in cancer cell lines that recapitulate APOBEC3-associated expression and mutation profiles observed in many human cancers. Our data show that BER components play a critical role in generating APOBEC3-associated mutations in breast and lymphoma human cancer cells. Finally, our results indicate important, but non-essential roles for APOBEC3A and APOBEC3B in generating different types of clustered mutations associated with APOBEC3 activities.

### Human cancer cell lines with active mutagenesis: models of APOBEC3 mutagenesis in cancer

To assess whether cell lines represent suitable models of APOBEC3 mutagenesis we compared APOBEC3-associated mutational signatures across DNA sequences of 780 widely used human cancer cell lines and 1,843 human cancers (Fig. 1b). The prevalence of the SBS2 and SBS13 in cell lines closely resembled their prevalence across the matching types of cancers, whereby cancers of breast, bladder, cervix and lung are among the most affected^4,5,30^. The appearance of the APOBEC3-associated signatures across human cell lines suggests that these signatures do not reflect a common mutational process associated with *in vitro* cultivation. Instead, APOBEC3-associated signatures in cell lines reflect traces of the exposures that in part occurred while the individual cell lineages were still evolving in vivo in cancer patients from which the cell lines were derived.

To determine the relative contributions of individual genes to generation of APOBEC3-associated signatures, we deleted a selection of candidate genes from two commonly used human breast cancer cell lines (BT-474 and MDA-MB-453), as well as two B cell lymphoma cell lines (BC-1 and JSC-1) (Fig. 1c,d; Extended Data Fig. 1; Extended Data Fig.). These cell lines naturally acquire APOBEC3-associated mutations over time^4^. Single-cell derived wild-type or knockout “parent” clones were subjected to long-term cultivation of 60-143 days corresponding to a timeframe over which mutation acquisition was investigated. Following this period, a further round of subcloning was carried out on the cell population from each of these parent clones. Multiple single-cell “daughter” clones were derived and shortly propagated to obtain DNA sufficient for analysis. In total, 136 individual parent and daughter clones were obtained and subjected to whole-genome sequencing (Table S1). This workflow enabled the detection of mutations unique to daughter clones thus identifying mutations acquired *de novo* over a defined period of *in vitro* propagation (Fig. 1d; Extended Data Fig. 3; Table S2; Table S3).

Examination of SBS profiles of the bulk cell lines revealed that BT-474, MDA-MB-453 and JSC-1 cell lines carried patterns of both SBS2 and SBS13, while BC-1 displayed only the SBS2 signature (Fig. 1e)^4^. *De novo* identification of mutational signatures from a total of 815,923 SBS discovered across 136 clones and 4 bulk cell line samples revealed evidence of six ongoing mutational processes (Fig. 1f; Table S4). Decomposition of these admixed patterns into previously identified SBS signatures revealed the presence of APOBEC-associated signatures SBS2 and SBS13^5^ (Table S4). SBS1 and SBS5, signatures of processes that operate continuously across most normal and cancer cells^5,31^, were also present (Table S4). Other identified signatures included SBS30, associated with inactivating mutations in the BER gene NTHL1^32^, and SBS8, SBS18 and SBS36, signatures of C>A mutations commonly attributed to oxidative stress in primary cancers and *in vitro* cultures^4,33–35^. The burdens of all mutational signatures were next quantified across individual wild-type and knockout cell line clones to investigate the contributions of candidate genes to acquisition of APOBEC-associated mutations.

### APOBEC3A drives acquisition of SBS2 and SBS13 in human cancer cells

As expected, ongoing generation of SBS2 and SBS13 was detectable in wild-type clones of all cell lines (Fig. 2a-l; Extended Data Fig. 4). APOBEC3-associated mutational burdens varied across individual daughter clones, consistent with previously reported episodic acquisition of these signatures in cancer cell lines (Fig. 2a-l; Extended Data Fig. 4; Table S4)^4^. This was most prominent in the BC-1 cell line, where for example, BC-1 daughter A.9 acquired 12,598 APOBEC3-associated SBS2 and SBS13 mutations in 108 days while a daughter A.10, which was propagated in parallel and derived from the same parent clone, exhibited only 1,807 of the respective mutations (Fig. 2k, Table S4). Analysis of cytosine mutations at APOBEC3A-preferred YT<u>C</u>A/YT<u>C</u>N and APOBEC3B-preferred RT<u>C</u>A/RT<u>C</u>N sequence contexts (Y=pyrimidine base, R=purine base, N=any base)^8^ revealed enrichment of the cytosine mutations in APOBEC3A-preferred contexts (Fig. 2f,h,j,l) across wild-type clones, corresponding to the enrichment of mutations in such contexts in most cancers^8^.

**Figure 2.**
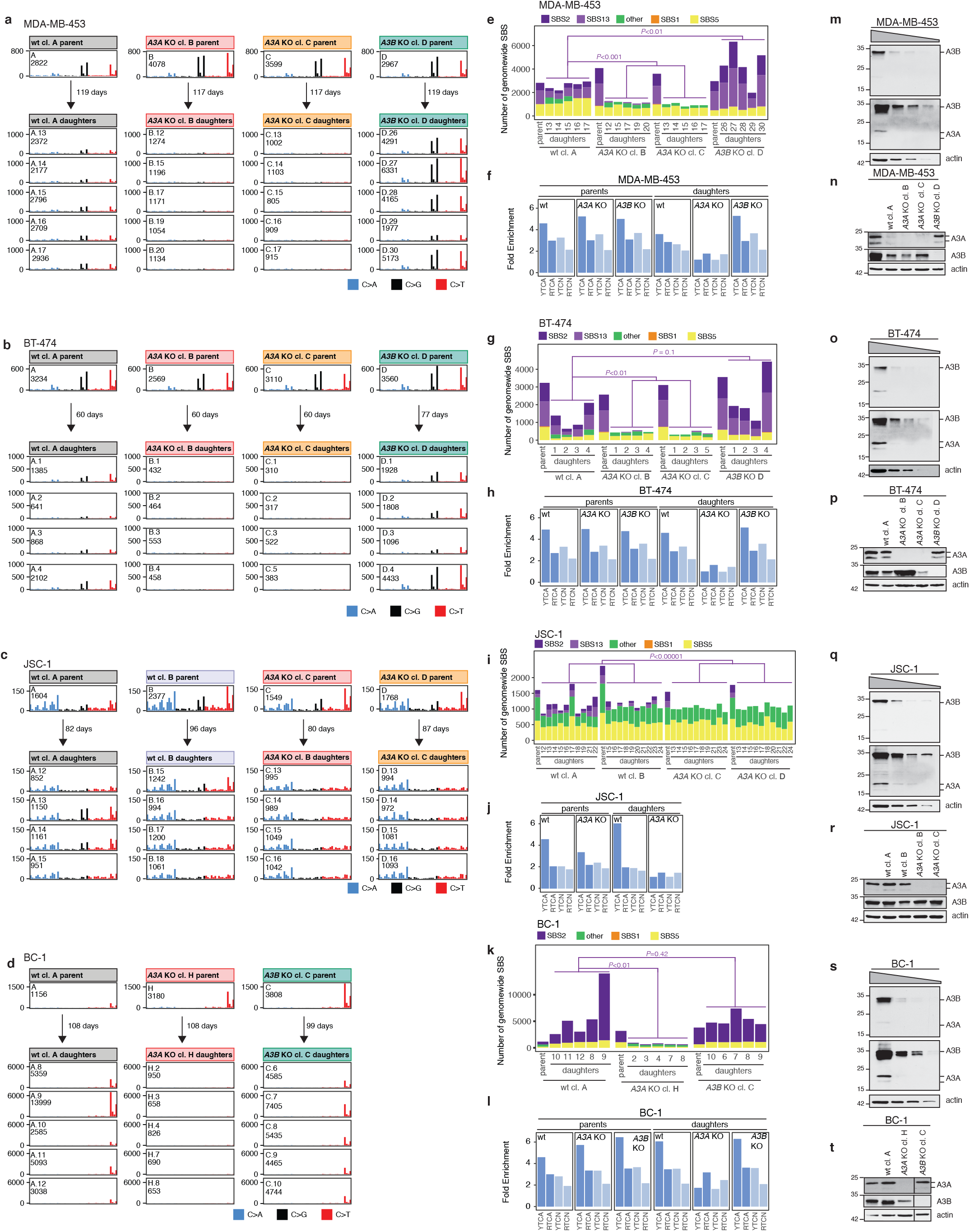
APOBEC3 deaminases drive acquisition of SBS2 and SBS13 in human cancer cells. **a-d)** Mutation acquisition in the indicated cell lines. Each panel is displayed according to the counts (y-axis) of genome-wide 48 cytosine base substitution classes denoted on horizontal axis, defined by the three color-coded cytosine base substitution types and 16 possible alphabetically ordered trinucleotide sequence contexts at which each base substitution type presents (order of cytosine substitution types follows standard representation, detailed in Extended Fig. 4). Arrows represent the number of days spanning the two subcloning events during which mutational acquisition was tracked, as in Fig. 1c. Additional JSC-1 clones shown in Extended data Fig. 5. **e-l)** Bars represent (e,g,i,k) genome-wide numbers of base substitutions attributed to discovered mutational signatures or (f,h,j,l) enrichment of cytosine mutations at APOBEC3B-preferred RT<u>C</u>A/N and APOBEC3A-preferred YT<u>C</u>A/N sequence contexts (R = purine base, Y = pyrimidine base, N = any base, mutated base is underlined) across mutational catalogues of annotated parent and daughter clones from denoted cell lines. *P* values were calculated by one-tailed Mann-Whitney *U* test to assess significant differences in SBS2 and SBS13 accumulation across cell lines. **m-t)** Immunoblotting with anti-APOBEC3 (04A04) and anti-actin antibodies using extracts (40 µg, 20 µg, 10 µg, and 5 µg) prepared from the indicated cell lines. Note that the anti-APOBEC3 antibody can detect both APOBEC3A and APOBEC3B (Extended Data Fig. 5a,b). Multiple exposures are shown to better depict APOBEC3A and APOBEC3B signals.

Consistent with widely reported observations of upregulation of *APOBEC3B* in breast and other cancer types^12,13,36^, all cell lines exhibited substantially elevated mRNA and protein levels of APOBEC3B relative to APOBEC3A (Fig. 2m,o,q,s; Extended Data Fig. 5a-g). Analyses across individual wild-type clones revealed that *APOBEC3A* and *APOBEC3B* expressions varied, but *APOBEC3B* was uniformly more abundant than the minimally expressed *APOBEC3A*. In line with its elevated expression levels, APOBEC3B represented the major cytidine deaminase activity directed against linear and hairpin probes in extracts prepared from MDA-MB-453 cells (Extended Data Fig. 5h-k). However, as reported before^15^, the presence of cellular RNA in extracts inhibited APOBEC3B activity, revealing that both APOBEC3A and APOBEC3B were enzymatically active against hairpin loop substrates in MDA-MB-453 cells (Extended Data Fig. 5l,m). In contrast to previous reports^15,16^, neither APOBEC3A nor APOBEC3B emerged as the dominant activity under these conditions. Deletion of each paralog elicited comparable losses in deaminase activity and removal of both *APOBEC3A* and *APOBEC3B* was required to eliminate deaminase activity. Thus, high expression levels and deaminase activity seemingly implicate APOBEC3B as the major mutator in all cancer cell lines analyzed here, while analyses of extended sequence contexts favor a role for APOBEC3A. These findings recapitulate widely reported findings that produced the ongoing debate regarding the relevance of each paralog in causing mutations in cancer^3,10^.

To test whether endogenous APOBEC3 activity represents an enzymatic source of cancer mutagenesis and delineate potential roles of candidate APOBEC3 paralogs, *APOBEC3A* and *APOBEC3B* were deleted by CRISPR-Cas9 gene targeting (Fig. 2n,p,r,t; Extended Data Fig. 1; Extended Data Fig. 5d-g). The expression levels of non-targeted APOBEC3 paralogs fluctuated across both wild-type and knockout clones, but were not systematically affected by gene targeting (Extended Data Fig. 5d-g). Despite low expression of *APOBEC3A* compared to *APOBEC3B* in all breast and lymphoma cell lines, and measurable activities from both enzymes upon DNA substrates *in vitro*, deletion of *APOBEC3A*, but not *APOBEC3B*, severely diminished SBS2 and SBS13 mutations in daughter clones isolated from knockout parent clones (Fig. 2a-l; Extended Data Fig. 4; Table S4). For example, daughter clones isolated from a wild-type MDA-MB-453 parent clone acquired, on average, 1049 ± 280 SBS2 and SBS13 mutations in 119 days while the daughter clones isolated from two of the MDA-MB-453 *APOBEC3A* knockout cell lines exhibited 45 ± 59 of the corresponding mutations over 117 days of culture (Fig. 2e). Similarly, *APOBEC3A* knockouts of BT-474 cells and both BC-1 and JSC-1 B cell lymphoma cell lines exhibited severely diminished accumulation of SBS2 and SBS13 mutations (Fig. 2g,i,k; Table S4). Although strongly diminished, APOBEC3-associated SBS2 and SBS13 mutations were not completely eliminated in many of the *APOBEC3A* knockout daughter clones from BT-474, MDA-MB-453 and BC-1 cell lines, indicating that additional APOBEC3 member(s) may be generating smaller burdens of mutations in these samples. Indeed, deletion of *APOBEC3A* was accompanied by a shift in the enrichment of mutations from APOBEC3A-preferred YTCN to APOBEC3B-preferred RTCN sequence contexts in daughter clones (Fig. 2f,h,l), suggesting that APOBEC3B may also cause mutations. Taken together, these experiments implicate APOBEC3A as the main driver of SBS2 and SBS13 in breast and B cell lymphoma lines and suggest that another APOBEC3 enzyme with a likely preference for RTCN motifs, such as APOBEC3B, may also contribute.

While deletion of *APOBEC3B* did not diminish overall mutational burdens, daughter clones isolated from the *APOBEC3B* knockout breast cancer cell line MDA-MB-453 exhibited significantly more SBS2 and SBS13 mutations than its wild-type counterparts (Fig. 2e; Table S4). This was not apparent in the BC-1 and BT-474 cell lines and could not be investigated in the JSC-1 cell line where *APOBEC3B* knockouts were not successfully established. Analyses of extended sequence contexts across all *APOBEC3B*-deleted clones revealed that the increased mutational burdens are enriched in APOBEC3A-preferred YTCN sequence contexts (Fig. 2f,h,j,l). The increase in mutations in the MDA-MB-453 cell line was reminiscent of the higher APOBEC3-associated mutational burdens observed in breast cancers that develop in carriers of a common germline deletion polymorphism that effectively deletes *APOBEC3B*^17,20^. The mechanisms underlying these observations remain unknown.

Burdens of SBS5 occasionally varied in clones from the MDA-MB-453 and BC-1 cell lines, albeit not as substantially as burdens of SBS2 and SBS13 (Fig. 2e,k; Table S4). SBS30, SBS8, SBS18 and SBS36 contributed small numbers of mutations compared to other signatures. The sums of mutations attributed to these signatures were thus represented together (‘other’) and fluctuated across individual clones due to mutational burdens that were underpowered for accurate quantification. (‘other’; Fig. 2e,g,i,k; Table S4).

### Base-excision repair plays a critical role in generation of APOBEC3 mutations in cancer

To assess the impact of BER on the generation of SBS2 and SBS13 in cancer cells (Fig. 1a), the uracil glycosylase *UNG* was deleted in BT-474 and MDA-MB-453 cells by CRISPR-Cas9 editing. SMUG1, which can occasionally substitute for UNG^37^, was removed from BT-474 cells. Successful gene targeting was confirmed by PCR and Sanger sequencing and loss of expression was verified by immunoblotting (Fig. 3a,b; Extended Data Fig. 2a). In contrast to wild-type clones from MDA-MB-453 and BT-474 cell lines, which exhibited both SBS2 and SBS13, daughters isolated from the *UNG* knockout clones exhibited exclusively SBS2 mutations (Fig. 3c-f; Table S4). This confirms that generation of transversion mutations in SBS13 depends on UNG-dependent uracil excision following APOBEC3-mediated cytosine deamination.

**Figure 3.**
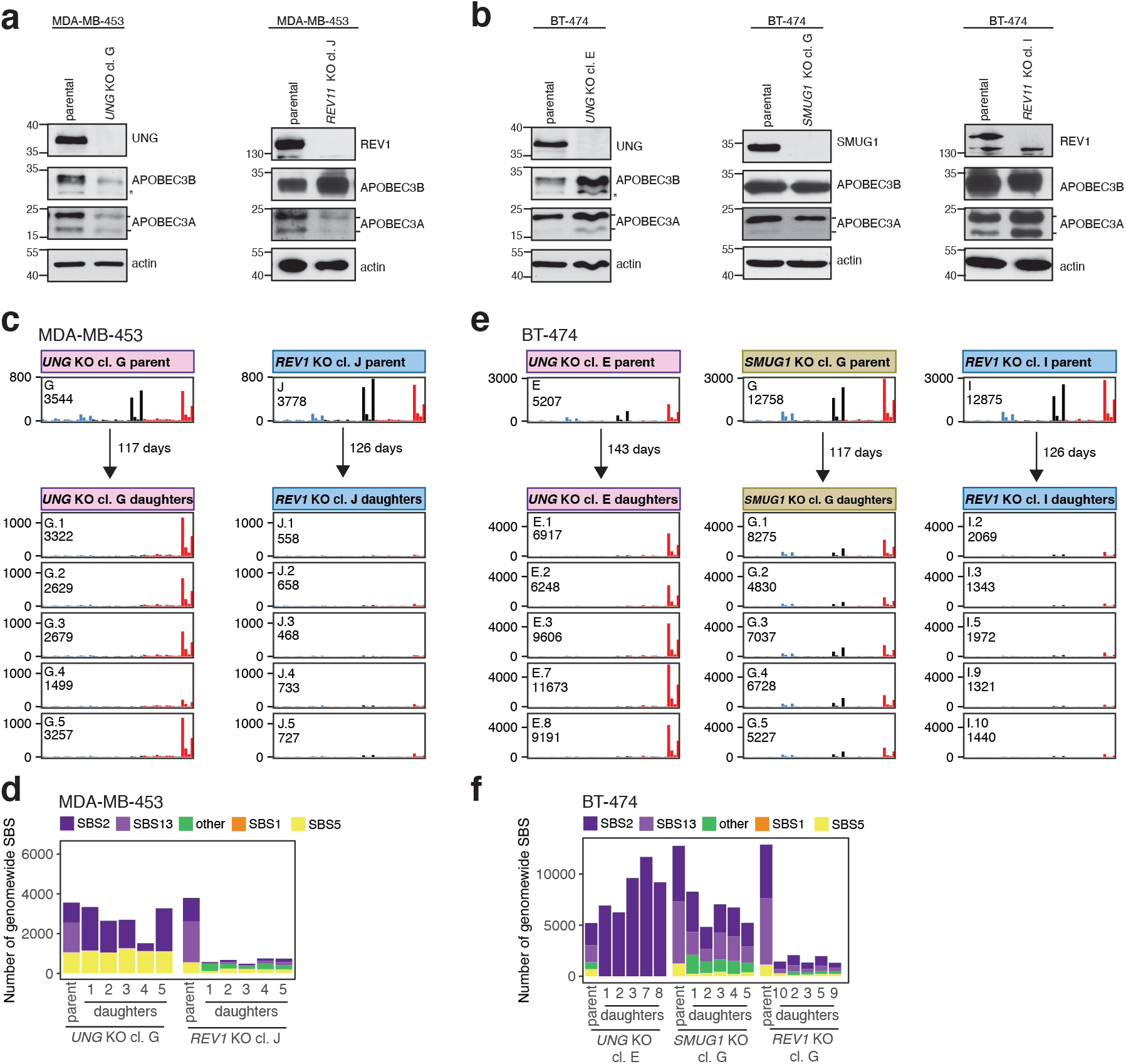
Base excision repair plays a critical role in generation of APOBEC3 mutations in cancer. **a-b)** Immunoblotting with anti-APOBEC3 (04A04), anti-UNG, anti-REV1, and anti-actin antibodies in the indicated cell lines. Note that the anti-APOBEC3A/B/G monoclonal detects long and short APOBEC3A isoforms. Asterisks mark nonspecific signals. **c**,**e** Mutation acquisition in the indicated cell lines. Each panel is displayed according to the counts (y-axis) of genome-wide 48 cytosine base substitution classes denoted on horizontal axis and defined by the indicated color-coded SBS types and 16 possible alphabetically ordered trinucleotide sequence contexts at which each mutation type presents (order of cytosine substitution types follows standard representation, detailed in Extended Fig. 4). Arrows represent the number of days spanning the two subcloning events during which mutational acquisition was tracked, as in Fig. 1e. **d**,**f)** Annotation of color-coded mutational signatures. Bars represent base substitutions attributed to mutational signatures in annotated clones.

Surprisingly, *UNG* deletion did elicit a major increase in the combined burden of SBS2 and SBS13 mutations in MDA-MB-453 cells, where *UNG* knockout clones were propagated for a similar number of days as wild-type clones (respectively, 117 and 119 days) (P=0.07, Mann-Whitney test; Fig. 2a,e; Fig. 3c,d; Table S4). Thus, most uracils generated by APOBEC3A base editing can be converted into C>T mutations by UNG-independent mechanisms.

Deletion of the nuclear uracil DNA glycosylase *SMUG1* did not affect the ability of BT-474 cells to acquire SBS2 and SBS13 (Fig. 3e,f; Table S4), indicating that SMUG1 is dispensable for the generation of SBS2 and SBS13. The observed dependency on UNG for the processing of APOBEC3A-generated uracils may derive from its ability to process both single-stranded and double-stranded DNA (dsDNA), while SMUG1 activity is essentially specific to dsDNA^38^.

Following uracil excision, replication across abasic sites by translesion synthesis (TLS) polymerases has been speculated to give rise to C>A and C>G transversions, as well as a portion of C>T mutations^39,40^. REV1 is proposed to form a scaffold for components of TLS during somatic hypermutation mediated by the AID APOBEC family member and to thus play a critical role in generation of a broad range of TLS-associated mutations^41^. To assess the contribution of TLS to generation of SBS2 and SBS13, *REV1* was targeted by CRISPR/Cas9 editing in breast cancer cell lines and loss of expression was verified by immunoblotting (Extended Data Fig. 2d-f). Consistent with the role of REV1 during AID-mediated somatic hypermutation^41,42^, this led to almost a 6-fold decrease in SBS2 and SBS13 in *REV1* knockout clones compared to wild-type clones in MDA-MB-453 cells (Fig. 3c-f; p=4.0×10-3, Mann-Whitney test) and more than a 4-fold decrease of the relevant signatures in *REV1* knockout clones compared to *UNG/SMUG1* knockout clones that were propagated for a similar number of days in BT-474 cells (both p=4.0×10-3, Mann-Whitney test) (Fig. 3c,d). These results suggest that REV1 plays a critical role in the generation of both SBS2 and SBS13. Substantial depletion of SBS2 signature mutations in *REV1*, but not *UNG* KOs, suggests that REV1 may have a key role in generation of C>T mutations that is independent of BER. Diminished SBS2 and SBS13 in the REV1 knockouts could not be attributed to perturbed growth or reduction in APOBEC3A levels (Fig. 3a, b; Extended Data Fig. 3). Nevertheless, we cannot exclude the possibility that APOBEC3A mutagenesis was synthetically lethal or selected against in *REV1* knockout cells.

Unlike SBS1, mutational burdens attributed to SBS5 were significantly depleted in *REV1* knockout cells of MDA-MB-453 cell lines (p=4.0×10-3, Mann-Whitney test). SBS5 has been attributed to an unknown process that is continuously operative across all tissues^31,43^ and its increased burdens in bladder cancers have been associated with mutations in the *ERCC2* gene encoding a DNA helicase that plays a central role in the NER pathway^43^. Our data suggests that REV1 may play a critical part in the underlying mutational process.

### APOBEC3 deaminases drive acquisition of *kataegis* and *omikli* mutations in human cancer cells

Most APOBEC3-associated mutations in examined clones were non-clustered (Fig. 4a). However, all cell lines acquired additional smaller numbers of clustered mutations, which commonly presented at the APOBEC3-associated cytosine mutations in TCN sequence contexts, including *kataegis* foci of densely clustered SBS mutations, *omikli* clusters of more sparsely distributed SBS mutations and doublet base substitutions (DBS) (Fig. 4a).

**Figure 4.**
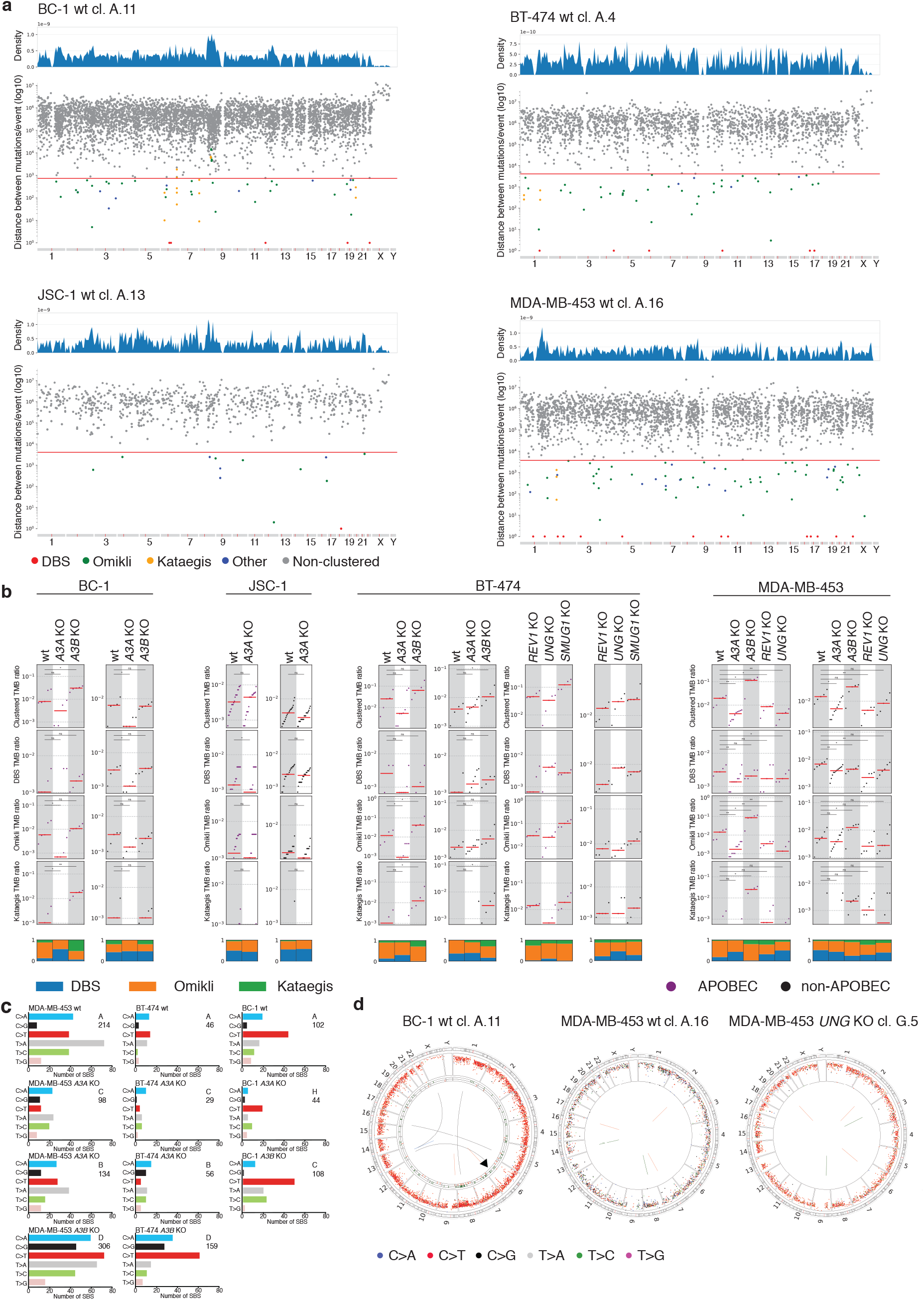
APOBEC3 deaminases drive acquisition of clustered mutations in human cancer cells. **a)** Rainfall plots of mutations acquired during the periods of defined *in vitro* growth in a selection of clones. Each dot represents a single base substitution, color-coded according to mutation-type (DBS = double-base substitution). The distances between mutations are plotted on the vertical axes on a log scale. The sample-dependent intermutation distance cutoffs for clustered mutations are shown as red lines, while regional corrections were performed to account for megabase heterogeneity of mutation rates. Mutation density plots are shown above each rainfall plot depicting the normalized mutation densities across the genome that were used for the regional corrections. **b)** Distribution of clustered APOBEC-like mutations (purple; cytosine mutations at TCN contexts) and all other mutations (non-APOBEC like; black), acquired *de novo* in daughter clones from designated cell lines and experiments. The total clustered tumor mutational burden (TMB) defined as mutations per megabase is further subclassified into the TMB of doublet-base substitutions, *omikli* associated events, and *kataegic* events, where each red bar reflects the median mutational burden for a given set of clones. A Mann-Whitney *U* test was performed for all statistical comparisons. Types of clustered events across each experiment are shown as bar-plots with each color proportionate to the events observed across all clones. **c)** Mutation spectra of clustered mutations in non-APOBEC-like contexts acquired *de novo* in designated clones. **d)** Circos plots depict mutations acquired *de novo* in denoted daughter clones. Color-coded SBS are plotted as dots in rainfall plots (log intermutation distance). Arrows point to examples of *kataegis*. Central lines indicate rearrangements (gray = translocations, green = tandem duplications, blue = inversions; orange = deletions).

Deletion of *APOBEC3A*, but not *APOBEC3B*, resulted in reduced burdens of *kataegis* foci and *omikli* clusters in BC-1, MDA-MB-453 and BT-474 cell lines at APOBEC3-like TCN sequence contexts (Fig. 4b). The *kataegis* losses were not detected in JSC-1 cells, which displayed minimal numbers of the relevant clusters. Indeed, consistent with the increased burden of genome-wide SBS2 and SBS13 observed in *APOBEC3B*-deleted clones from MDA-MB-453 (see section *‘APOBEC3A drives acquisition of SBS2 and SBS13 in human cancer cells’*), there was an elevated number of APOBEC3-like *kataegis* foci in *APOBEC3B* knockout clones from all cell lines and APOBEC3-like *omikli* was increased in *APOBEC3B* knockout clones from the breast cancer cell lines (Fig. 4b). Neither APOBEC3A nor APOBEC3B were required for generation of *kataegis* and *omikli*, as both were occasionally observed in the relevant knockout daughters. Taken together these data indicate that APOBEC3A is the main driver of APOBEC-like *kataegis* and *omikli*, but suggest that additional mutators, such as APOBEC3B, may play a minor role as previously proposed^44^.

Unexpectedly, loss of *APOBEC3A* also caused a reduction in clustered mutations occurring outside of APOBEC3-like sequence contexts in BC-1 and MDA-MB-453 cells, while deletion of *APOBEC3B* led to their modest increase in breast cancer cell lines (Fig. 4b,c). These SBS primarily consisted of C>T transitions, consistent with the possibility that they may derive, in part, from non-canonical APOBEC3A base editing at exposed regions of ssDNA.

*Kataegis* foci often co-localize with rearrangements in primary cancers, a phenomenon attributed to APOBEC3 attacks on ssDNA exposed during the resection phase of homologous recombination-mediated DNA double-strand break repair^9,45^. A separate explanation proposes that APOBEC3-induced deamination may precede the dsDNA breaks, if ssDNA breaks generated upon UNG-mediated uracil excision represent the initiating lesions for formation of subsequent dsDNA breaks^9^. In line with the latter proposal, burdens of APOBEC-like clustered mutations were reduced in *UNG* knockout clones, compared to wild-type clones from the MDA-MB-453 cell line. However, UNG was not essential for *kataegis* in MDA-MB-453 and BT-474 cell lines (Fig. 4b), nor in the BC-1 cell line where UNG expression is attenuated^4^. Additionally, there were several examples of *kataegis* foci that appeared to occur independently of any proximal rearrangements in cell line clones (Fig. 4d). These data suggest that *kataegis* can occur independently of APOBEC3-initiated DNA cleavage likely at spontaneous DNA breaks or uncoupled DNA replication forks^46^. However, all clones acquired small numbers of rearrangements and we cannot exclude the possibility that initiating DNA double strand breaks were successfully repaired as cell lineages harboring chromosome rearrangements may have been selected against during *in vitro* propagation (Extended Data Fig. 6). Finally, in line with REV1 contributing to a broader spectrum of SBS mutations (Fig 3.c-f), including non-clustered signatures SBS5 and APOBEC-associated SBS2 and SBS13, deletion of *REV1* in MDA-MB-453 cells resulted in reduced mutational burdens of clustered mutations occurring both within and outside of the APOBEC3-like sequence contexts.

## DISCUSSION

This study provides the first direct evidence for a hypothesis formulated in 2002^29^, which speculated that APOBEC3 cytidine deaminases may represent potent mutators in human cancer cells. The data establish APOBEC3A as the main driver of highly prevalent genome-wide and clustered *kataegis* APOBEC3-associated mutational signatures, in breast and B cell lymphoma cancer cells.

APOBEC3-associated mutational signatures are enriched at YTCN sequence contexts in the majority of individual human cancers and cancer types^8,12,13^. Our finding that APOBEC3A accounts for most APOBEC-associated mutations at YTCN sequence contexts in human cancer cells strongly indicates that APOBEC3A drives acquisition of the large majority of all APOBEC-associated mutations observed in cancer genomes, as has been speculated before based on observations in yeast^8^. All the cancer cell lines analyzed in this study, where APOBEC3A is the predominant driver of the relevant mutations, possess high levels of APOBEC3B expression relative to APOBEC3A, an observation that was previously used to nominate APOBEC3B as the major mutator in cancer^12,13,36^. Furthermore, despite APOBEC3A being the predominant mutator, activities of APOBEC3A and APOBEC3B were similar in *in vitro* deamination assays that have commonly been used as substitute readouts of mutagenesis by individual enzymes^12,13^. Thus, the data shows that increased expression and deamination activities of individual APOBEC members may not always translate into active mutagenesis. These findings caution against the widespread use of such readouts as sole substitute measures of active mutagenesis by APOBEC3 deaminases, which resulted in distinct predictions regarding APOBEC members as predominant mutators in cancer^12,13,15,47^. The direct measurements of mutagenic activities of APOBEC3A and APOBEC3B enzymes in human cancer cell line genomes used here represent the strongest available support that mutagenesis by APOBEC3A, and not APOBEC3B, represents the major source of some of the most prevalent mutational signatures in human cancer. Recent work, largely based on correlations between individual APOBEC3 expression levels and deamination activities, has implicated distinct APOBEC3 members as drivers of targeted therapy resistance in lung cancers^48,49^. Our results call for the use of more direct measures of APOBEC3 activity to delineate the role of individual APOBEC3 enzymes in cancer genome evolution.

The presented data cannot exclude the possibility that APOBEC3B or other APOBEC family members cause mutations. Indeed, although SBS2 and SBS13 mutations were substantially depleted in *APOBEC3A* knockout clones, they were not completely eliminated, suggesting that other enzymes may play a minor role. It is also conceivable that stable *APOBEC3B* expression across longer time periods than those analyzed in this study may result in a more substantial contribution to SBS2 and SBS13 mutational burden. Our study also cannot account for potential cell-type specific differences that may impact APOBEC3 activity. In a smaller proportion of cancers that are enriched in APOBEC3B-preferred RTCNA motifs, most prominently in lung adenocarcinomas^8^, APOBEC3B may be a more relevant mutator than APOBEC3A. Contributions of individual APOBEC3 family members to different stages of cancer evolution will require further investigation. Finally, our data implicate UNG and REV1, and thus BER, to the generation of APOBEC3-induced non-clustered signatures SBS2 and SBS13, as well as clustered *kataegis* and *omikli* events in cancer cell genomes.

Experimental confirmation of APOBEC3 deaminases as mutators in human cancer cells and identification of APOBEC3A as the main generator of widespread mutations in cancer marks a critical advance in pursuing the proposed therapeutic interventions based on modulating the generation of the associated SBS signatures^21–28^ and in investigating the origins of APOBEC3-associated mutations in cancer. Our data suggest that uncovering the factors that drive misregulation of APOBEC3A will be critical to identify the sources of many mutations in cancer and that modulation of mutagenic activities by APOBEC3A may offer avenues for the proposed therapeutic interventions^17,27,50^.

## Data availability statement

All sequencing data pertaining to this project have been deposited in the European Nucleotide Archive database with the accession number ERP108795. All the other data supporting the findings of this study are available within the article and its supplementary information files and from the corresponding authors upon reasonable request. Source data will be provided at publication.

## Code availability

The code used in this study is available at the Wellcome Sanger Institute GitHub page (https://github.com/cancerit) and was published before.

## Acknowledgements

We thank members of the Maciejowski lab for critical reading of this manuscript and Lisa Mohr for help with formatting. Work was supported by a Cancer Grand Challenges Mutographs team award funded by Cancer Research UK (C98/A24032) and by the Wellcome grant reference 206194. Work in J.M.’s laboratory is supported by the NCI (R00CA212290; P30 CA008748), the Pew Charitable Trusts, the V Foundation, the Starr Cancer Consortium, the Emerald Foundation, and the Geoffrey Beene and Ludwig Centers at MSKCC. M.P. is supported by the European Molecular Biology Organization (EMBO) Long-Term Fellowship (ALTF 760-2019). This work was also supported by the US National Institute of Health grants R01ES030993-01A1 and R01ES032547-01 to L.B.A..

## Additional information

Correspondence and requests for materials should be addressed to M.P. and J.M.

## Author Contributions

M.P. and J.M. conceived and designed the study. M.P. and J.M. wrote the manuscript with contributions from M.R.S. K.C., A.D., P.V.M, J.M. performed the experiments. M.P. analyzed the genomics data. E.N.B. and L.B.A performed analyses on clustered mutations. All authors approved the final manuscript.

## Methods

### Data Reporting

No statistical methods were used to predetermine sample size. The investigators were not blinded to allocation during experiments and outcome assessment.

## Cell Culture

MDA-MB-453, BT-474, JSC-1, and BC-1 cell lines were acquired from the cryopreserved aliquots of cell lines sourced previously from collaborators or public repositories and extensively characterized as part of the Genomics of Drug Sensitivity in Cancer (GDSC)^1,2^ and COSMIC Cell Line projects^3,4^. Bulk cell lines were genotyped by SNP and STR profiling, as part of the COSMIC Cell Line Project (https://cancer.sanger.ac.uk/cell_lines) and individual clones obtained here were genotyped (Fluidigm) to confirm their accurate identities. MCF10A cells were from Maria Jasin’s lab (MSKCC).

Annexin V staining was performed using the annexin V Apoptosis detection kit (BD Biosciences) according to the manufacturer’s instructions.

### Generation of Knockout Cell Lines

10^6^ cells were electroporated using the Lonza 4D-Nucleofector X Unit (MDA-MB-453) or Lonza Nucleofector 2b Device (BT-474, BC-1, JSC-1) using programs DK-100 (MDA-MB-453), X-001 (BT-474), or T-001 (BC-1, JSC-1) in buffer SF + 18% supplement (MDA-MB-453) or 80% Solution 1 (125 mM Na_2_HPO_4_•7H_2_O, 12.5 mM KCl, acetic acid to pH=7.75) and 20% Solution 2 (55 mM MgCl_2_) (BT-474, BC-1, JSC-1) and 9 µg (UNG, SMUG1, REV1) or 10 µg (A3A, A3B) of pU6-sgRNA_CBh-Cas9-T2A-mCherry plasmid DNA (Table S5). mCherry positive cells were single-cell sorted into 96-well plates by FACS using FACSAria (BD Biosciences).

### Knockout Screening and Validation by PCR

#### CRISPR KO Clone Screening

Genomic DNA isolated using a Genomic DNA Isolation Kit (Zymo Research; cat. ZD3025). Purified genomic DNA for CRISPR/Cas9 knockout screens was amplified using Touchdown PCR. Each PCR reaction consisted of: 7.4 µL ddH_2_O, 1.25 µL 10× PCR buffer (166 mM NH_4_SO_4_, 670 mM Tris base pH 8.8, 67 mM MgCl2, 100 mM β-mercaptoethanol), 1.5 µL 10 mM dNTPs, 0.75 µL DMSO, 0.25 µL forward and reverse primers (10 µM each), 0.1 µL Platinum Taq DNA Polymerase (Invitrogen; 10966083), and 1 µL genomic DNA. Primer sequences are listed in Table S5.

#### PCR for Sanger Sequencing

PCR reactions for Sanger Sequencing were performed using the Invitrogen Platinum Taq DNA Polymerase (Invitrogen; 10966083) protocol. 25 ng of genomic DNA was used for each reaction. Primer sequences are listed in Table S5. DNA from PCR reactions was purified from agarose gels using the Invitrogen PureLink Quick Gel Extraction Kit (Invitrogen; K210012). Gel-purified DNA was cloned using the TOPO TA Cloning Kit for Sequencing (Invitrogen; 450030) and colonies were selected for sequencing (Genewiz).

### RNA Isolation and Quantitative PCR

RNA was isolated using a *Quick* -RNA Miniprep Kit (Zymo Research; R1054). RNA was quantified and converted to cDNA using the SuperScript IV First-Strand Synthesis System (Invitrogen; 18091050). cDNA synthesis reactions were performed using 2 µL of 50 ng/µL random hexamers, 2 µL of 10 mM dNTPs, 4 µg RNA, and DEPC-treated water to a volume of 26 µL. The mixture was heated at 65°C for 5 minutes, then cooled on ice for 5 minutes. Primers, probes, and cycling conditions were adopted from published methods^5^. Primer sequences are listed in Table S5.

### Immunoblotting

Cells were lysed in RIPA buffer (150 mM NaCl, 50 mM Tris-HCl pH 8.0, 1% NP-40, 0.5% sodium deoxycholate, 0.1% SDS, Pierce Protease Inhibitor Tablet, EDTA free) or sample buffer (125 mM Tris-HCl pH 6.8, 1 M β-mercaptoethanol, 4% SDS, 20% glycerol, 0.02% bromophenol blue). Quantification of RIPA extracts was performed using the Thermo Scientific Pierce BCA Protein Assay kit. Protein transfer was performed via wet transfer using 1× Towbin buffer (25 mM Tris, 192 mM glycine, 0.01% SDS, 20% methanol) and nitrocellulose membrane. Blocking was performed in 5% milk in 1× TBST (19 mM Tris, 137 mM NaCl, 2.7 mM KCl, and 0.1% Tween-20) for 1h at room temperature (RT). The following antibodies were diluted in 1% milk in 1× TBST: anti-APOBEC3A/B/G and anti-APOBEC3A (see below; WB 1:500), anti-APOBEC3B (Abcam; ab184990; WB 1:500), anti-REV1 (Santa Cruz; sc-393022, WB 1:500), anti-SMUG1 (Abcam; ab192240; WB 1:1,000), anti-UNG (abcam; ab109214; WB 1:1,000), anti-GFP (Santa Cruz; sc-9996; WB 1:1,000), anti-β-actin (Abcam; ab8224; WB 1:3,000), anti-β-actin (Abcam, ab8227; WB 1:3,000); anti-Mouse IgG HRP (Thermo Fisher Scientific; 31432; 1:10,000), anti-Rabbit IgG HRP (SouthernBiotech; 6441-05; 1:10,000).

### APOBEC3 monoclonal antibody generation

Residues 1-29 (N1-term) or 13-43 (N2-term) from APOBEC3A and residues 354-382 (C-term) from APOBEC3B and were used to create three peptide immunogens (EZBiolab). Five mice were given three injections using Keyhole-Limpet-Hemocyanin (KLH)-conjugated peptides over the course of 12 weeks (MSKCC Antibody and Bioresource Core). Test bleeds from the mice were screened for anti-APOBEC3A titers by ELISA against APOBEC3A peptides conjugated to BSA. Mice showing positive anti-APOBEC3A immune responses were selected for final immunization boost before their spleens were harvested for B-cell isolation and hybridoma production. Hybridoma fusions of myeloma (SP2/IL6) cells and viable splenocytes from the selected mice were performed by MSKCC Antibody and Bioresource Core. Cell supernatants were screened by APOBEC3A ELISA. The strongest positive hybridoma pools were subcloned by limiting dilution to generate monoclonal hybridoma cell lines. Hybridomas 04A04 and 01D05 were expanded then grown in 1% FBS medium. This medium was clarified by centrifugation and then passed over a Protein G column (04A04) or Protein A column (01D05) to bind mAb. The resulting mAb was eluted in PBS (04A04) or 100 mM NaCitrate pH 6.0, 150 mM NaCl buffer (01D05).

### *In vitro* DNA deaminase activity assay

Deamination activity assays were performed as described^6^. Briefly, 1 million cells were pelleted and lysed in buffer (25 mM HEPES, 150 mM NaCl, 1 mM EDTA, 10% glycerol, 0.5% Triton-X, 1× protease inhibitor), sheared through a 28 ½-gauge syringe, then cleared by centrifugation at 13,000 × g for 10 minutes at 4°C. Deaminase reactions (16.5 µl cell extracts with 2 µl UDG buffer (NEB), 0.5 µl RNase A (20 mg/ml), 1 µl 1 µM probe (linear = 5’IRD800/ATTATTATTATTATTATTATTTCATTTATTTATTTATTTA or hairpin = 5’IRD800/ATTATTATTATTGCAAGCTGTTCAGCTTGCTGAATTTATT), and 0.3 µl UDG (NEB)) were incubated at 37°C for 2 hours followed by addition of 2 µl 1M NaOH and 15 minutes at 95°C to cleave abasic sites. Reactions were then neutralized with 2 µl 1 M HCl, terminated by adding 20 µl urea sample buffer (90% formamide + EDTA) and separated on a pre-warmed 15% acrylamide/urea gel in 1× TBE buffer at 60°C for 70 minutes at 100V to monitor DNA cleavage. Gels were imaged by Odyssey Infrared Imaging System (Li-COR) and quantified via ImageJ.

### Comparison of APOBEC3-associated mutational signatures in cell line with cancer data

Annotations of mutational signatures across 1,001 human cancer cell lines and 2,710 cancers from multiple cancer types were published previously^3^. Where possible, we matched cancer and cell line cancer classes as detailed in Table S1. Eventually, 780 cell lines and 1843 cancers from matching types were used in analyses presented in Fig. 1b. Individual classes and samples per class used are listed in Table S1, while the signature annotation was published previously^3^ and downloaded here.

### Whole-genome Sequencing

Genomic DNA was extracted from a total of 136 individual clones using the DNeasy Blood and Tissue Kit (QIAGEN) and quantified with Biotium Accuclear Ultra high sensitivity dsDNA Quantitative kit using Mosquito LV liquid platform, Bravo WS and BMG FLUOstar Omega plate reader. Samples were diluted to 200ng/120µl using Tecan liquid handling platform, sheared to 450bp using a Covaris LE220 instrument and purified using Agencourt AMPure XP SPRI beads on Agilent Bravo WS. Library construction (ER, A-tailing and ligation) was performed using ‘NEB Ultra II custom kit’ on an Agilent Bravo WS automation system. PCR was set up using Agilent Bravo WS automation system, KapaHiFi Hot start mix and IDT 96 iPCR tag barcodes or unique dual indexes (UDI, Ilumina). PCR included 6 standard cycles: 1) 95°C 5 mins; 2) 98°C 30 s; 3) 65°C 30 s; 4) 72°C 1 min; 5) cycle from 2, 5 more times; 6) 72°C 10 mins. Post-PCR plates were purified with Agencourt AMPure XP SPRI beads on Beckman BioMek NX96 liquid handling platform. Libraries were quantified with Biotium Accuclear Ultra high sensitivity dsDNA Quantitative kit using Mosquito LV liquid handling platform, Bravo WS and BMG FLUOstar Omega plate reader, pooled in equimolar amounts on a Beckman BioMek NX-8 liquid handling platform and normalized to 2.8 nM ready for cluster generation on a c-BOT. Pooled samples were loaded on the Illumina Hiseq X platform using 150 PE run lengths and sequenced to approximately 30× coverage, as detailed in Table S1. Sequencing reads were aligned to the reference human genome (GRCh37) using Burrows-Wheeler Alignment (BWA)-MEM (https://github.com/cancerit/PCAP-core). Unmapped, non-uniquely mapped reads and duplicate reads were excluded from further analyses.

### Mutation calling

Somatic single base substitutions (SBS) were discovered using CaVEMan (https://github.com/cancerit/cgpCaVEManWrapper)^7^, with major and minor copy number options set to, respectively, 5 and 2, to maximize discovery sensitivity. Rearrangements were identified with the BRASS algorithm (https://github.com/cancerit/BRASS). Sequences of the corresponding parent clones were used as reference genomes to discover mutations in individual daughter clones, whereas a sequence from an unrelated normal human genome^3^ was used as a reference to discover mutations in parent clones. Individual comparisons are outlined in Table S1. Mutations shared between parent clones (see below) were used to derive proxies for the mutational catalogues of bulk cell lines (Fig. 1e). Rearrangements were retained only if identified as absent from the reference sequences by BRASS. SBS discovered with CaVEMan were filtered over the two additional steps: first, to remove the low-quality loci and, second, to ensure that the mutational catalogues from daughter clones retained exclusively mutations acquired during the relevant *in vitro* periods spanning the two cloning events and that the mutational catalogues from parent clones retained predominantly mutations acquired prior to the examined *in vitro* periods. Individual comparisons performed and the numbers of mutations removed with individual filters are in Table S2.

First, only SBS flagged as ‘PASS’ by Caveman when analyzed across the panel of 98 unmatched normal samples (https://github.com/cancerit/cgpCaVEManWrapper)^7^ were considered, removing large proportions of mapping and sequencing artefacts, as well as the common germline variation^7^. Four post-hoc filters were applied to ‘PASS’ variants to further remove sequencing and mapping artifacts that occur with XTEN and BWA-mem-aligned data and to ensure that the mutation loci were sufficiently covered in the reference sequences. ‘PASS’ mutations were removed if (Filter 1; Table S2) the median alignment score (ASMD) of mutation-reporting reads was less or equal to 140; if (Filter 2; Table S2) the mutation locus had the clipping index (CLPM) greater than 0; if (Filter 3; Table S2) the mutation locus was covered by 20 or less reads in the reference samples used in comparisons; and if (Filter 4; Table S2) less than two sequencing reads of opposite directions reported the mutation.

Second, we genotyped all mutation loci which passed the filters above across all available clones from the matching cell lines. We used cgpVAF (https://github.com/cancerit/vafCorrect) to count the number of mutant and wild type reads across individual clones. Mutations from each parent or daughter clone that were found at cumulative VAF of >5% across >10% of clones from other parental lineages were removed (Filter 5, Table S2). Mutations presenting at clones from other parental lineages below these cut-offs were determined false-positive calls upon manual inspection of individual reads and were thus retained. In mutational catalogues from parent clones, this step served to remove the majority of the germline mutations and a smaller proportion of mutations shared between parent clones, thus retaining predominantly somatic mutations acquired in individual parent cell lineages prior to the examined *in vitro* periods spanning the two cloning events. In mutational catalogues from daughter clones, the filter served to remove mutations which presented across clones from other parental lineages and were thus likely acquired before examined *in vitro* periods, but were not captured in the corresponding reference sequences. The likely pre-existent germline and somatic mutations that were shared between the related parent clones were accumulated into mutational catalogues of bulk cell lines (Fig. 1e). The percentages of mutations removed with this filter also represent the upper-level estimates of the remaining false-positive *de novo* SBS calls in mutational catalogues from daughter clones, which may not have been captured in the reference sequences and may have been designated as *de novo*. Such mutations may have been removed by filtering against other parental lineages, but their estimated proportions do not affect results and are generally minor (median ∼2.5%; per-sample estimates in Table S2). Finally, while this filter removes most of the germline and the pre-existing variation, a smaller proportion of the removed mutations may have arisen independently across multiple parental lineages at the hairpin loci that are hotspots for APOBEC3-associated mutagenesis^8^.

### Validation of parent-daughter allocations

Genotyping of remaining mutation loci across all clones revealed that, rarely, a large proportion of mutations absent from the parent clones was shared between some or all daughters (e.g. Extended Data Fig.7c, BC-1_C lineage daughter clones). To exclude the possibility that high proportions of shared mutations stem from allocations of the relevant daughters to the wrong parents, we confirmed the presence of the expected CRISPR-edits in genome sequences from all such daughters (not shown) and we confirmed that such shared mutations were absent from all other clones from individual cell lines Extended Data Fig.7a-d). This originally revealed a swap between two lineages and a couple of clones from JSC-1 cell line (not shown), which are annotated in Table S1 and resolved in all data representations (including Extended Data Fig.7a-d). A few clones that exhibited a higher level of sharedness were not resolved in this way, (e.g. daughters from BC-1_C lineage; BC-1_H.3 and BC-1_H.8; see Extended Data Fig.7c). To exclude the possibility of clone cross-contaminations, in which case VAF of shared mutations would be lower than VAFs of other clonal mutations in some clones, we confirmed that the VAF distributions of shared mutations followed those of other clonal mutations (not shown).

In the absence of sample swaps and putative contaminations, rare instances where high proportions of clonal mutations were shared between the related daughters and absent from their corresponding parents indicate that the corresponding daughters were most likely established from the common subclone that arose during the cultivation of the parent clone, after its DNA was already extracted.

### Validation of clonal sample origins

To ensure that samples were clonal and single–cell-derived, we examined proportions of the variant-reporting reads (equivalent to variant allele fraction, VAF) at the mutation loci (Extended Data Fig. 7e). Consistent with the polyploid background of most cell lines under investigation^3^, VAF distributions often deviated from the average of ∼50% expected for clonal heterozygous somatic mutations occurring in a diploid genome. The largely unimodal VAF distributions confirmed the clonal origins of the majority of the samples. In occasions where bimodal VAF distributions were observed, at least one of the peaks followed the VAF distribution of all the other related clones, indicating that the other peak originates from mutations acquired subclonally. Such instances were overall rare and most common in the BC-1 cell line.

### Sequence context-based classification of single base substitutions

SigProfilerMatrixGenerator (python v.1.1; https://github.com/AlexandrovLab/SigProfilerMatrixGenerator)^9^ was used to categorize SBSs into three separate sequence-context based classifications. The algorithm allocates each SBS to (1) one of the 6-class categories (C>A, C>G, C>T, T>A, T>C and T>G) in which the mutated base is represented by the pyrimidine of the base pair; (2) to one of the 96-class categories (in which each of 6-class mutation types is further split into 16 subcategories baked on the flanking 5′ and 3′ bases); (3) and to one of the 1,536-class categories (in which each of 6-class mutation types is further split into 256 subcategories based on two flanking bases 5′ and 3′ to the mutated base). Relevant outputs are in table Table S3.

### Enrichment of APOBEC3-associated mutations at target motifs

Once SBSs were allocated to their sequence context classes as described, whereby the mutated base is represented by the pyrimidine base of the base pair, C>T and C>G base substitutions at TCN (N is any mutation) contexts which brand APOBEC3-associated SBS2 and SBS13 signatures were classified as ‘APOBEC3-associated’, whereas C>T and C>G substitutions at other contexts were classified as ‘OTHER’. C>A substitutions were excluded because some of the C>A mutations have been attributed to both APOBEC3 mutagenesis, as well as other mutational processes commonly arising during *in vitro* cell cultivation^3^. Enrichment of ‘APOBEC3-associated’ mutations was then investigated in the specific pentanucleotide motifs^10^ across all clones.

### Enrichment of APOBEC3-associated mutations at trinucleotide and pentanucleotide motifs

Enrichment of APOBEC3-associated mutations was compared across the pentanucleotide motifs that were previously associated with APOBEC3A (YTCN and YTCA, where Y is a pyrimidine base) and APOBEC3B activities (RTCN and RTCA, where R is a purine base) in yeast overexpression systems^10^. Relevant APOBEC3-associated trinucleotide and pentanucletide sequence motifs were quantified with sequence_utils (v.1.1.0, https://github.com/cancerit/sequence_utils/releases/tag/1.1.0; (https://github.com/cancerit/sequence_utils/wikisequence-context-of-regions-processed-by-caveman) across human autosomal chromosomes (GRCh37) and by excluding the regions not considered by the CaVEMan algorithm in detecting SBS. Middle base pair of each reference pentanucleotide sequence was considered a putative mutation target and the surrounding sequence context was extracted by using the DNA strand belonging to the pyrimidine base of the target base-pair. A total of 96 possible trinucleotide and 512 pentanucleotide contexts were quantified across both DNA strands (e.g. AGT trinucleotide is reported as ACT; AAGCA pentanucleotide is reported as TGCTT; middle ‘target’ bases underlined). Enrichment of ‘APOBEC3-associated’ mutations at the pentanucleotide motifs of interest was calculated as described previously^3,10^. For example, to calculate enrichment of ‘APOBEC3-associated’ mutations at RTCN sites the following was used:

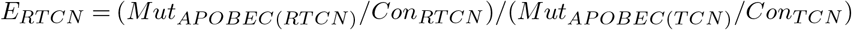

Mut_APOBEC(TCN)_ is the total number of ‘APOBEC3-associated’ mutations (C>G and C>T mutations at T<u>C</u>N contexts) in autosomal chromosomes; Mut_APOBEC(RTCN)_ is the sum of ‘APOBEC3-associated’ mutations at RT<u>C</u>N contexts in autosomal chromosomes; whereas Con_TCN_ and Con_RTCN_ represent the total number of T<u>C</u>N and RT<u>C</u>N contexts available among the regions considered by Caveman when calling mutations across the autosomal chromosomes. As described, both DNA strands are considered, but the mutation types and target motifs are reported based on the strand of the pyrimidine base of the target base pair.

### Mutational signatures analysis

Mutational signatures analyses were performed using the SigProfilerExtractor tool (v. 1.0.17; https://github.com/AlexandrovLab/SigProfilerExtractor)^11^, which is a method based on nonnegative matrix factorization (NMF) for *de novo* extraction of mutational signatures from a given matrix of SBS types. SBS were classified into 96 classes based on their trinucleotide sequence contexts (see ‘Sequence context-based classification of single base substitutions’). The tool was used over 500 iterations to identify profiles of mutational signatures operative across a total of 815,923 genome-wide mutations identified across 4 bulk cell lines and their corresponding 136 daughter and parent clones. Mutational signatures were extracted *de novo* and mapped to the known COSMIC Mutational Signatures of cleaner patterns derived from more powered cancer datasets (v3, https://cancer.sanger.ac.uk/cosmic/signatures; see Table S4). Activities of identified COSMIC mutational signatures were quantified in each clone as part of the factorization of the input 96-SBS channel matrices, whereby numbers of SBS mutations belonging to each signature were quantified in the genome of each sample. The relevant outputs from SigProfilerExtractor are in Table S4 and include profiles of *de novo* extracted signatures, metrics related to mapping of *de novo* signatures to COSMIC signature profiles and per-sample activity estimations. Statistical comparisons across clones were performed using a one-tailed Mann-Whitney *U* test.

### Identification of clustered mutations

To detect clustered single base substitutions, a sample-dependent inter-mutational distance (IMD) cutoff was derived, which is unlikely to occur by chance given the mutational pattern and mutational burden of each clone. To derive a background model reflecting the distribution of mutations that one would expect to observe by chance, SigProfilerSimulator (v1.1.2) was used to randomly simulate the mutations in each clone across the genome^12^. Specifically, the model was generated to maintain the +/- 1bp sequence context for each substitution, the strand coordination including the transcribed or untranscribed strand within genic regions^9^ and the total number of mutations across each chromosome for a given sample. All single base substitutions were randomly simulated 100 times and used to calculate the sample-dependent IMD cutoff so that 90% of mutations below this threshold were clustered with respect to the simulated model (i.e., not occurring by chance with a q-value<0.01). Further, the heterogeneity in mutations rates across the genome and the variances in clonality or copy-number were considered by correcting for mutation rich regions present in 10Mb-sized windows and by using a threshold for the difference in variant allele frequencies between subsequent substitutions in a clustered event (variant allele frequency difference<0.10). Subsequently, the clustered mutations were subclassified into specific categories of events: *(i)* doublet substitutions; two adjacent mutations with consistent variant allele frequencies; *(ii)* extended multi-base substitutions; previously termed *omikli* events^13^ that reflect any two mutational events greater than 1bp and less than the sample-dependent IMD cutoff with consistent variant allele frequencies; *(iii)* large mutational events; previously termed kataegi^14^ with three or more mutational events greater than 1bp and less than the sample-dependent IMD cutoff with consistent variant allele frequencies. Lastly, statistical comparisons across clones were performed using a Mann-Whitney *U* test.

