## Extended Data Figures for "The APOBEC3A deaminase drives episodic mutagenesis in cancer cells"

Extended Data Fig. 1. Petljak et al.

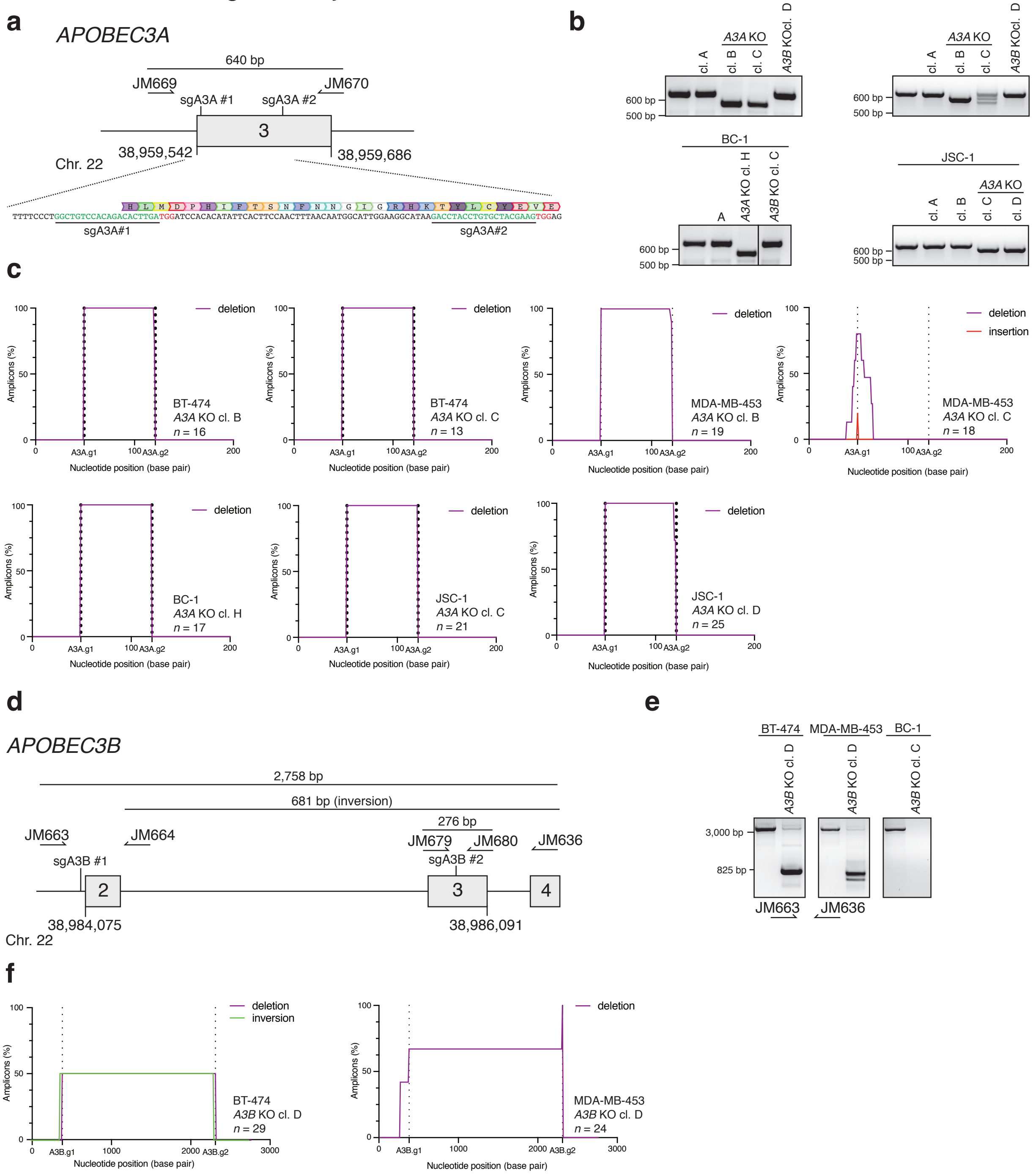


**Extended Data Figure 1. Generation of *APOBEC3A* and *APOBEC3B* knockout cell line clones. a)** Schematic of APOBEC3A locus. Position of exon 3, targeting sgRNAs (sgA3A 1 and sgA3A 2) and primers for PCR screening (JM669 and JM670) are indicated. **b)** PCR amplicons generated using primers JM669 and JM670 and genomic DNA templates prepared from the indicated cell lines. **c)** Plots depict a percentage of sequenced amplicons generated as in b) that contain deletions (purple) or insertions (red) at the indicated positions. **d)** Schematic of the *APOBEC3B* locus. Position of exons 2-4, targeting sgRNAs (sgA3B 1 and sgA3B 2) and primers for PCR screening (JM663 and JM636) are indicated. **e)** PCR amplicons generated using primers JM663 and JM636 and genomic DNA templates prepared from the indicated cell lines. **f)** Plots depict a percentage of sequenced amplicons generated as in e) that contain deletions (purple) and inversions (green).

Extended Data Fig. 2. Petljak et al.

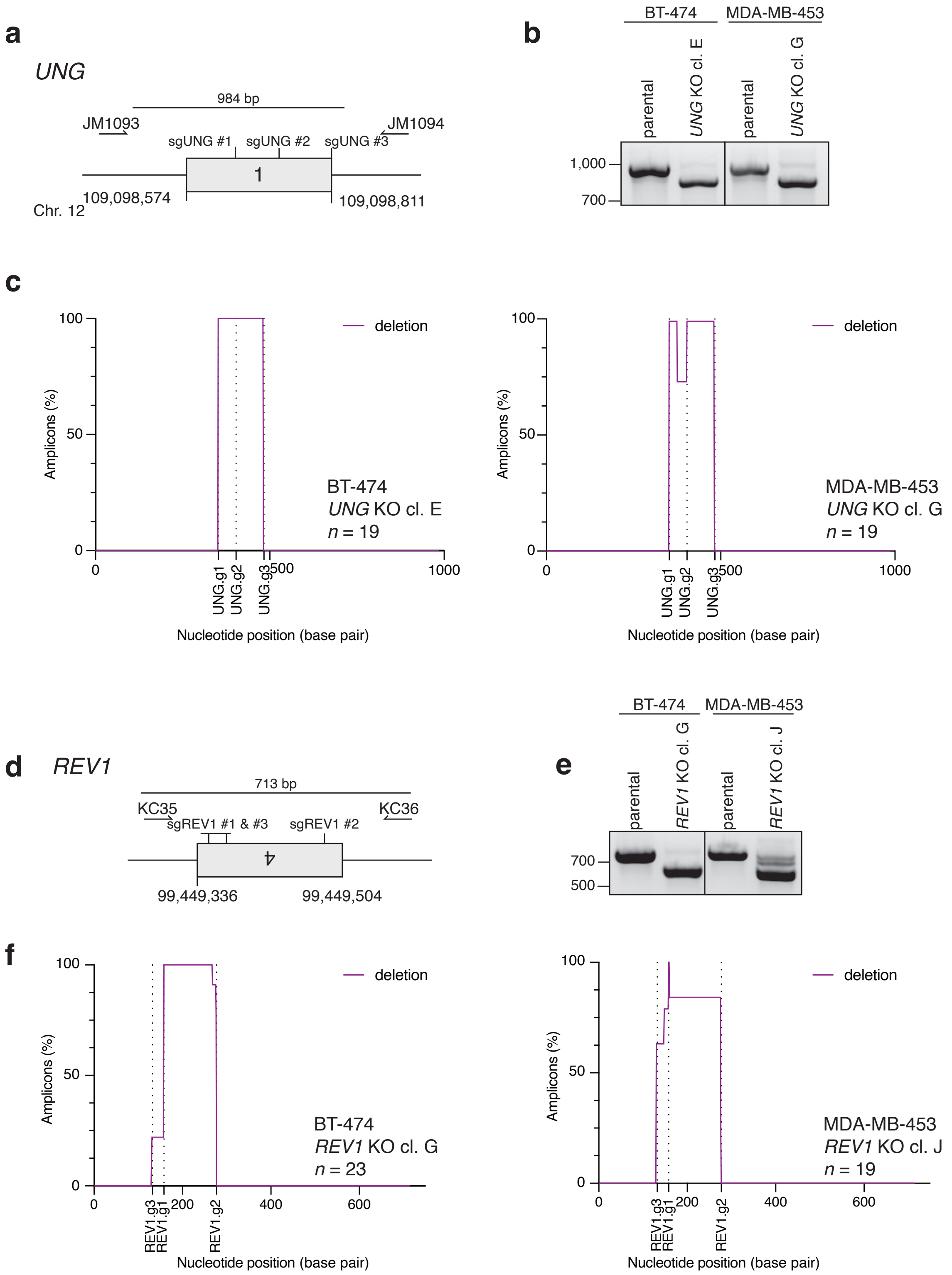


**Extended Data Figure 2. Generation of *REV1* and *UNG* knockout cell line clones. a)** Schematic of *UNG* locus. Position of exon 1, targeting sgRNAs (sgUNG 1 and sgUNG 2) and primers for PCR screening (JM1093 and JM1094) are indicated. **b)** PCR amplicons generated using primers JM1093 and JM1094 and genomic DNA templates prepared from the indicated cell lines. **c)** Plots depict a percentage of sequenced amplicons generated as in b) that contain deletions (purple). **d)** Schematic of *REV1* locus. Position of exon 4, targeting sgRNAs (sgREV1 1 and sgREV1 2), and primers for PCR screening (KC35 and KC36) are indicated. **e)** PCR amplicons generated using primers KC35 and KC36 and genomic DNA templates prepared from the indicated cell lines. **f)** Plots depict a percentage of sequenced amplicons generated as in d) that contain deletions (purple).

Extended Data Fig. 3. Petljak et al.

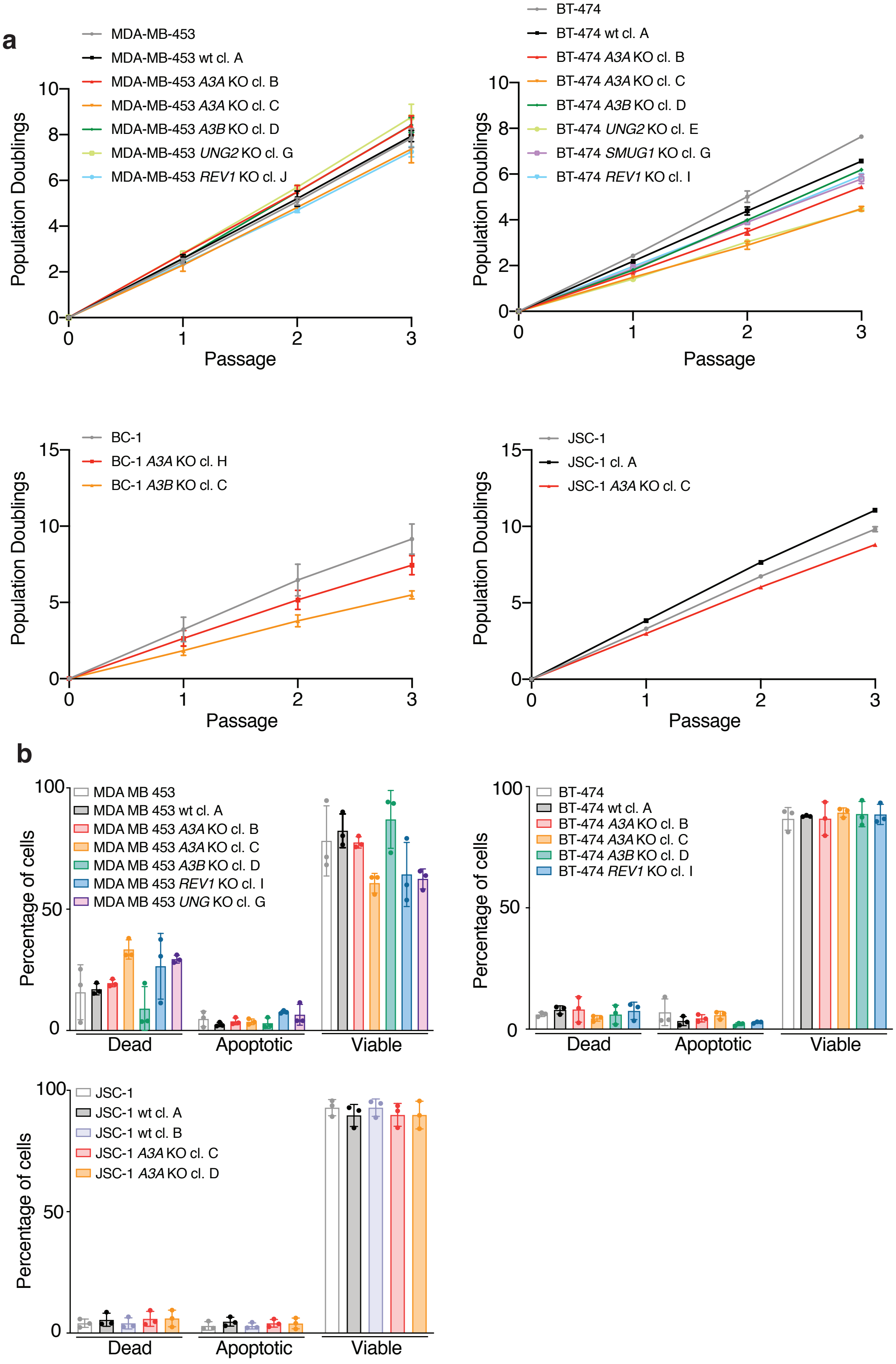


**Extended Data Figure 3.** (Caption next page.)

**Extended Data Figure 3.** (Previous page) **Population doubling and apoptosis in targeted cancer cell lines. a)** Population doubling measures over successive passages from the indicated cell lines. Mean and s.d. are from $n = 3$ independent biological replicates are shown. **b)** Plot showing percentages of apoptotic, necrotic, or living cells as indicated by propidium iodide and annexin V staining. Bars represent mean and s.d. from at least two independent experiments.

Extended Data Fig. 4. Petljak et al.

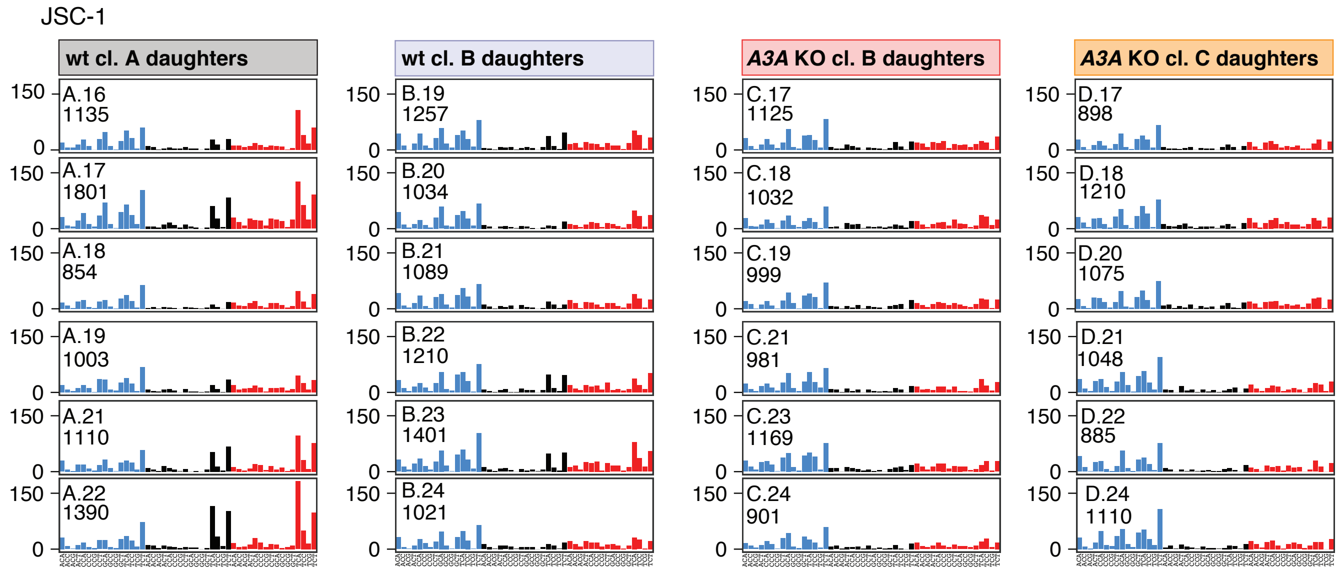


**Extended Data Figure 4. APOBEC3 deaminases drive acquisition of SBS2 and SBS13 in JSC-1 cell line.** Additional clones obtained from JSC-1 experiments presented as in Fig 2k.

Extended Data Fig. 5. Petljak et al.

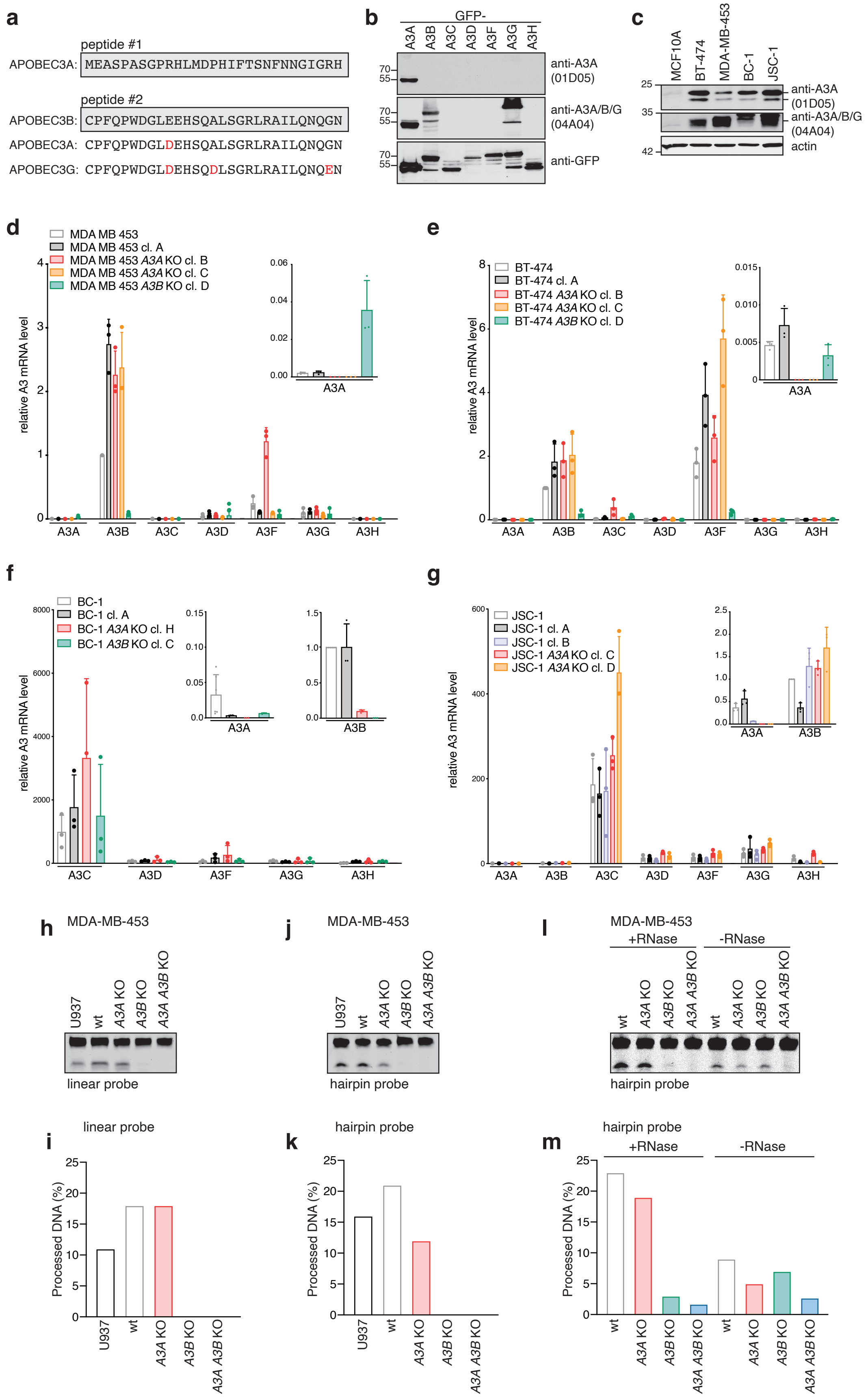


**Extended Data Figure 5.** (Caption next page.)

**Extended Data Figure 5.** (Previous page) **Expression and deamination activities of APOBEC3A and APOBEC3B across cell line clones. a)** Peptides used to generate anti-APOBEC3 (04A04) and anti-APOBEC3A (01D05) mouse monoclonal antibodies. **b)** Immunoblotting with anti-APOBEC3A (01D05), anti-APOBEC3 (04A04), and anti-GFP antibodies in extracts prepared from HEK293T cells transfected with the indicated GFP-APOBEC3 constructs. **c)** Immunoblotting with anti-APOBEC3A (01D05), anti-APOBEC3 (04A04), and anti-actin antibodies in the indicated cell lines. **d-g)** Normalized *APOBEC3* mRNA levels in the indicated cell lines based on qPCR. The mean and s.d. of $n = 3$ independent biological replicates are shown. **h,j,l)** Cytidine deaminase activity in the indicated cell lines measured against a **h)** linear probe after RNase treatment, **j)** hairpin probe after RNase treatment, and **l)** hairpin probe after vehicle or RNase treatment. **i,k,m)** Measurement of the percentage of processed DNA as in h,j,l).

### Extended Data Fig. 6. Petljak et al.

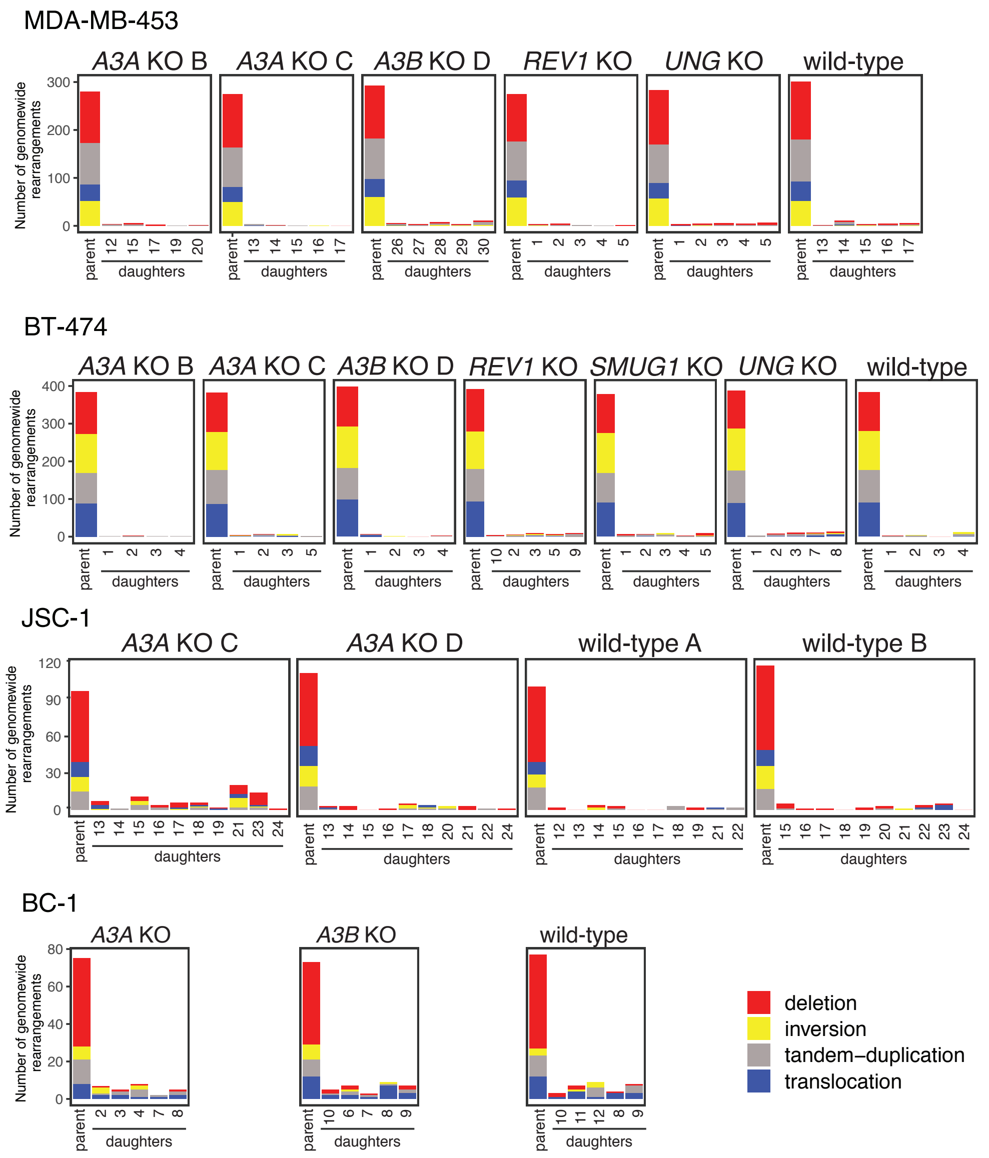


**Extended Data Figure 6. Analysis of chromosome rearrangements across cell line clones.** Plots showing numbers of color-coded rearrangement types detected genome-wide in the indicated cell line clones.

Extended Data Fig. 7. Petljak et al.

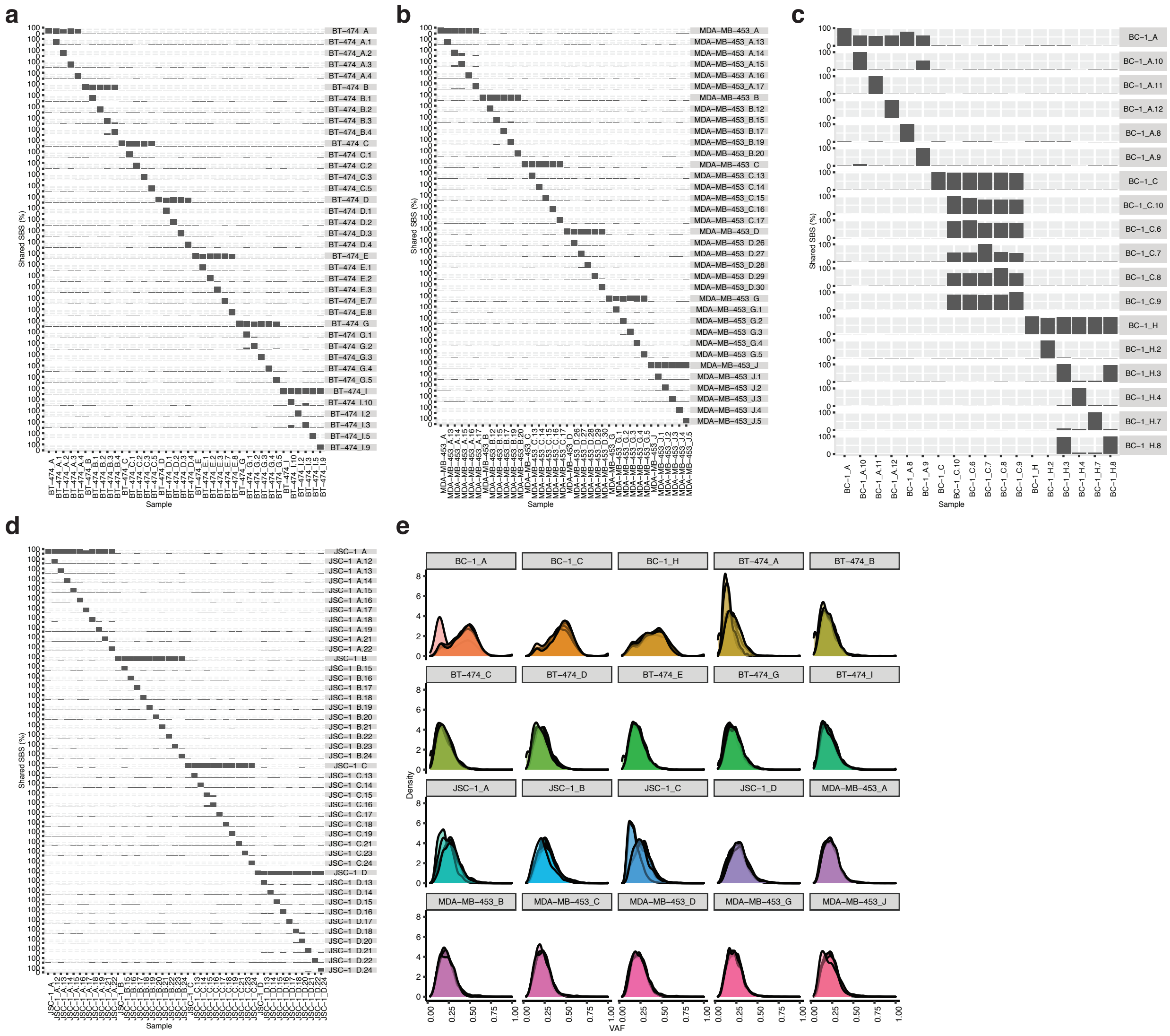


**Extended Data Figure 7. Distribution and variant allele fractions of mutations identified across all clones. a-d)** Left vertical axis depicts percentage of SBS mutations from **a)** BT-474, **b)** MDA-MB-453, **c)** BC-1 and **d)** JSC-1 cell line clones indicated on the right vertical axis that had been identified by across the related clones indicated on the horizontal axis upon genotyping of individual mutations (see Methods). **e)** Distributions of variant alleles fractions (VAFs) of mutations identified in individual parent and daughter clones from the experiments indicated on top. VAF peaks can sometimes deviate from 50%, expected for clonal heterozygous somatic mutations in a diploid genome, because cancer cell lines are often polyploid and heterozygous copy number changes across the genome can further modulate the VAF distribution. Bimodal distributions and subclonal peaks in wild-type clones from BC-1 cell line likely arose due to subclonal evolution of the relevant clones (Methods).
